# Subthalamic deep brain stimulation and chronic dopaminergic therapy enhance free-choice seeking in patients with Parkinson’s disease

**DOI:** 10.1101/2023.01.04.522610

**Authors:** David Bendetowicz, Gizem Temiz, Nicolas Tempier, Elodie Hainque, Marie-Laure Welter, Virginie Czernecki, Brian Lau, Carine Karachi, Jérôme Munuera

## Abstract

**Background:** Humans prefer making choices freely, even when they don’t maximize future outcomes, suggesting free-choice is intrinsically rewarding. However, whether reward-related brain networks influence choice preference remains unclear. In Parkinson’s disease (PD), value-based decision impairments are well-documented, but mechanisms underlying intrinsically motivated behavior are poorly understood. This study investigates how the dopaminergic and basal ganglia systems encode intrinsic reward in PD.

**Methods:** We designed a decision-making task dissociating free-choice’s intrinsic value from extrinsic reward. Twenty PD patients with subthalamic deep brain stimulation (STN-DBS) and twentyfive on dopamine (DA) therapy performed the task ON and OFF their treatments. Their performances were compared to twenty age-matched healthy controls. To explore neural mechanisms, we analyzed DBS active contacts, modeled the volume of tissue activated, and examined cortico-subthalamic connectivity using high-resolution diffusion MRI.

**Results:** PD patients OFF STN-DBS exhibited reduced free-choice preference, which increased when STN-DBS was ON, particularly in risky choices. This effect correlated with the recruitment of the right medial prefrontal cortex (mPFC). DA therapy did not modulate free-choice preference acutely, but higher chronic DA levels correlated with increased free-choice preference.

**Conclusions:** Our findings suggest STN-DBS enhances free-choice preference via the right mPFC-STN network, while chronic DA therapy amplifies free-choice sensitivity. This implies that free-choice preference is influenced by mPFC modulation, increasing impulsivity toward risky choices, and dopamine’s role in enhancing sensitivity to both extrinsic and intrinsic rewards.

## Introduction

Decisions are often described as a balancing act between rewards and punishments. However, rational decision-making models based on this balance frequently fail to accurately predict human behavior (1). Recent suggestions indicate that decisions can be influenced by value attribution that is not solely tied to extrinsic outcomes but also relies on intrinsic motivation. This concept is crucial in understanding behaviors driven by incentives that do not conform to economic optimization or can be easily explained by fulfilling basic needs such as hunger, thirst, sex, or pain avoidance (1–3). For example, multiple species show a preference for opportunities to choose, even when these choices do not provide increased extrinsic rewards (4–6). Similarly, humans tend to prefer having options to collect favorite items rather than being forced to make a selection, even if this choice incurs greater costs (7,8). Although the desire to choose may seem economically irrational, opportunities for choice can offer greater control over the environment, potentially triggering rewarding signals (3,8).

The dopaminergic (DA) networks and the basal ganglia system are implicated in decisions influenced by extrinsic rewards and punishments (9–11). These systems may also play a central role in processing intrinsic rewards (1,12). For instance, it has been shown that midbrain DA neurons in non-human primates encode the inferred value of cues, providing other forms of intrinsic rewards, such as information about future outcomes, without impacting the rate of external reward consumption (13). In humans, striatal activation increases when participants can freely choose different items, regardless of the magnitude of their associated extrinsic rewards or even when they are aware that free-choice is available. (14–16).

Impairments in value-based decisions are observed in pathological conditions such as Parkinson’s disease (PD), in which DA cell loss leads to profound basal ganglia dysfunction. Patients with PD exhibit impaired value-based learning, which can be enhanced through DA therapy (17–19). DA therapy has the potential to shift outcome value predictions to improve learning from rewards while minimizing the effects of adverse outcomes(17,18,20). Additionally, it has been shown that even though deep brain stimulation (DBS) targeting the subthalamic nucleus (STN) significantly diminishes patients’ motor fluctuations, it also alters decision-making, resulting in impulsive choices (21–24). This impulsivity may stem from the modulation of the prefrontal hyperdirect pathway, facilitating quick decision processes through response inhibition of sub-optimal behaviors (17-20), depending on the anatomical and functional organization of the STN and the positioning of the electrodes (25–30). While the posterior regions of the STN relate to motor functions, most anterior areas are associated with non-motor processes (26,27,30,31). However, although these manipulations have provided valuable insights into the roles of DA and prefronto-subthalamic networks in decision-making, they primarily emphasize external consequences (32,33).

To deepen our understanding of the cortico-subthalamic and DA systems’ contributions to intrinsic motivation, we designed an experiment in which PD patients express their intrinsic preference for collecting extrinsic rewards through either a free or forced choice option. We investigated the role of the cortico-subthalamic and DA systems in choice preferences by manipulating STN-DBS on one hand and DA therapy on the other. By altering the risk of losing rewards associated with free-choice, we explore how risk-seeking attitudes and their associated neural systems can modulate the value of intrinsic reward when it competes with the maximization of extrinsic reward intake.

## Methods and Materials

### Participants

We determined the sample size based on previous studies (8,16). We recruited 45 patients through the neurology and neurosurgery departments at the Pitié-Salpêtrière Hospital (Paris, France). DOPA patients (n=25, 7 females) received DA therapy only and were assessed during a follow-up visit to evaluate eligibility for DBS surgery, encompassing a comprehensive neuropsychological evaluation and an acute levodopa challenge test. DBS patients (n=20, 9 females) were assessed during a routine follow-up visit (delay after surgery: 15 months +/-5.3 Standard Deviation [SD]). For the comparative analysis, we enrolled twenty age and gender-matched, healthy controls (HC).

Inclusion criteria and details of stimulation devices and parameters of DBS patients are provided in Supplementary Methods and Supplementary Table 1.

**Table 1.** Statistical comparison of demographic data in the three groups of participants.

| Patient groups | DBS (n=20) | DOPA (n=25) | Healthy controls | Kruskall-Wallis or Wilcoxon's test |
| --- | --- | --- | --- | --- |
|  | Mean (SD) | Mean (SD) | Mean (SD) | (p value) |
| Age (years) | 54.2 (7.8) | 56.8 (8.6) | 54.2 (7.7) | K=2.33 (0.312) |
| MoCA (/30) | 27.5 (1.5) | 28.2 (1.4) | 28.1 (1.2) | K=3.41 (0.182) |
| Duration (years) | 12.3 (3.3) | 10.4 (4.3) | - | W=319.5 (0.113) |
| MDS-UPDRS III (OFF) | 47.7 (16.3) | 43.5 (14.5) | - | W=291 (0.355) |
| **MDS-UPDRS III (ON)** | **16.6 (9.9)\*** | **10.0 (7.3)** |  | **W=360.5 (0.012)** |
| **LEDD (mg)** | **405 (224)** | **1355 (525)** | - | **W=497 (<0.0001)** |

\* ON-DBS was assessed by keeping patients off dopamine replacement therapy
SD: Standard Deviation

The study was approved by the local ethics committee (CEEI/IRB n.21-810), and informed consent was given following the Declaration of Helsinki ethical rules.

### Task description

We developed a two-stage task using the Psychophysics Toolbox (http://psychtoolbox.org/) running on MATLAB (Natick, Massachusetts: The MathWorks Inc) that was previously described in a previous study (8). For each trial, during the first stage, two image targets were displayed on the screen, representing the *free* or *forced* options, and participants were required to make an initial selection. In the second stage, two different image targets were presented. In the *free* option, participants could select either of the two targets in the second stage, whereas, in the *forced* option, they had to select the one predetermined by the computer. During this forced condition, the computer consistently selected the same target, and the selection of the other image was disabled. The reward outcome was displayed after the second-stage target was selected (Figure 1).

**Figure 1.**
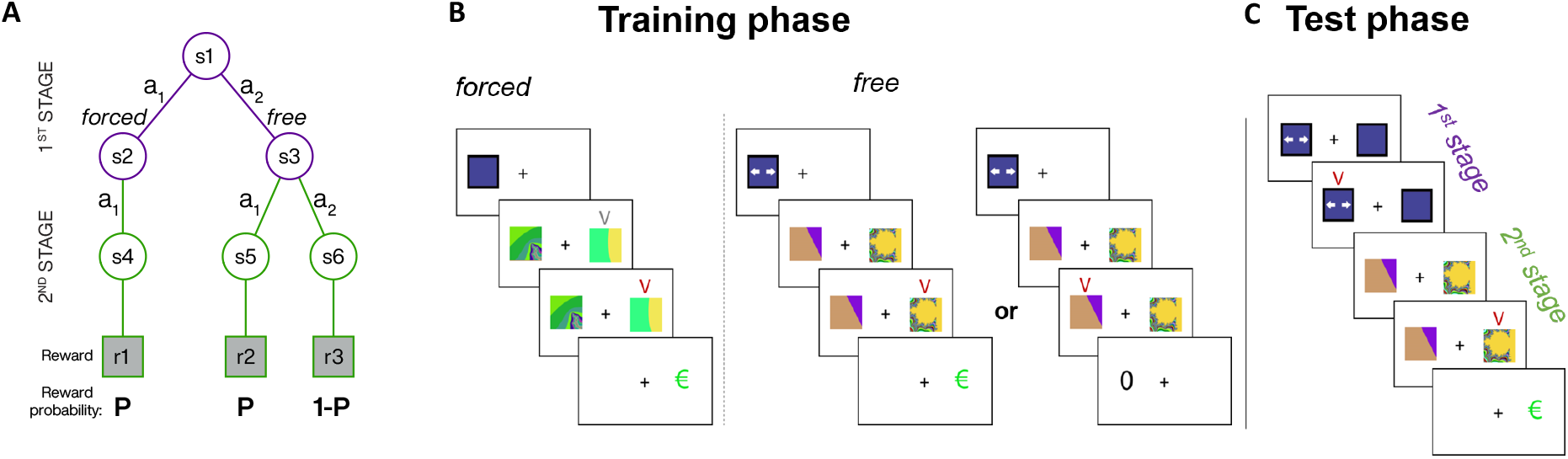
Two-stage task structure. (A) State diagram illustrating the six possible states (s), actions (a) and associated extrinsic reward probabilities (e.g., P = 0.5, 0.75 or 1 for blocks 1 to 3, respectively); s2 and s3 were represented by two different 1^st^-stage targets (e.g., colored squares with or without arrows for *free* and *forced* trials, respectively) and s4 to s6 were associated to three different 2^nd^-stage targets (fractals). (B) Illustration of typical trials during the training phase. Participants learned the reward contingencies (green euro symbol) associated with the second-stage targets in the *forced* and *free* options for each probability block. When training for forced choice, participants had to select the target indicated by a V-shaped grey symbol. Participants could freely select one of the two targets when training for free-choice. The V-shaped red symbol indicates the participant’s selection after the keypress. (C) Schematic illustration of a trial during the test phase. In the first stage, participants had to select whether to perform the trial in the *free* or *forced* option.

Reward contingencies were predetermined across three different blocks of 48 trials each. In one block, reward delivery was stochastic, with all second-stage targets associated with a reward probability (P) of 0.5. In a second block, the reward probabilities were unbalanced, with one second-stage target set at P=0.75 and the other at P=0.25. In the final block, reward delivery was deterministic: one second-stage target was rewarded at P=1 while the other was never rewarded (P=0). Importantly, the computer consistently selected the most rewarded target during the *forced* option. Thus, the reward contingencies were identical between the *free* and *forced* options, although each first- or second-stage image varied across all blocks. Before each testing phase, participants underwent a preliminary training phase to learn the contingencies between the distinct targets and their associated reward probabilities. They were clearly informed that the stimuli and their associated reward probabilities were identical across the training and testing phases.

### Procedure

DOPA patients performed the task during an acute levodopa challenge test (34). OFF sessions took place after discontinuation of levodopa treatment for 12 hours and DA agonist treatments for 72 hours. ON sessions took place the same day, 45 minutes after taking the levodopa equivalent dose of their usual morning treatment plus 50 mg of levodopa.

DBS patients performed the task after a complete wash-out of DA therapy for both sessions. They performed the task ON-DBS, then OFF sessions started at least an hour after the stimulation had stopped.

To account for any test-retest effect, HC performed the task twice with a one-hour interval between the two sessions. There was no difference in performance between the two sessions (p=0.707) and nor interaction with reward probability (p=0.792; Supplementary Figure 1).

### Statistical analyses

Generalized linear mixed models (GLMMs) were employed for binary dependent variables (first- and second-stage choices). We examined intrinsic preference for free-choice by evaluating the frequency of participants’ selection of the *free* option during the initial stage of the task. Linear mixed models (LMMs) were used for reaction times (RTs). All models included all sessions (DBS-ON/OFF, DOPA-ON/OFF, HC) as fixed effects, with HC serving as the reference level, along with its interaction with the average obtained reward. RT models also included the selected option (*free* or *forced*) and all interactions. Random effects included intercepts and slopes of the obtained reward by patients in all models. The models were fitted using the lme4 package in RStudio, applying Wald’s z statistic and corresponding P-values (R Core Team, https://www.R-project.org). Post-hoc statistics were adjusted for multiple comparisons using the Tukey method. Results are presented using estimated marginal means. Two-sample Wilcoxon tests were employed for demographic statistics.

### Tractography analysis

We assessed structural connectivity between the prefrontal cortex and the STN area in each patient treated with DBS using volume of tissue activated (VTAs) modeling and a high-definition normative local PD connectome (35,36). This normative connectome was created using multi-shell diffusion-weighted imaging (DWI) data from a local cohort of 33 PD patients candidates for DBS (26). We used the Lead-DBS V2 pipeline for electrode reconstruction, VTA modelling and co-registration between the pre-and post-surgery brain imagery. The spatial normalization was done in the Montreal Neurological Institute reference space, and the coherence between the individual and the normative space was verified patient by patient (37). The full image processing pipeline using Lead-DBS is available on the supplementary methods.

The cortical connectivity based on the Brodmann areas (BA) segmentation was extracted from the whole brain normative tractogram. For each extracted cortico-VTA connectivity, corresponding track density (TD) maps were created using SIFT2 weights for statistical analysis. These 3D density maps reflect the estimated weight of streamlines within the voxel (38). Voxelwise analysis was performed in the TD maps using permutation analysis of linear models (PALM) from FSL (https://fsl.fmrib.ox.ac.uk/fsl/fslwiki/PALM). We compared the TD maps of DBS patients with a difference in free-choice preference between ON and OFF sessions with those whose preference remained the same by median splitting the DBS group based on ON-OFF difference. To focus on the specificity of non-motor processes and restrain the number of voxelwise comparisons, the analysis was performed on a cortical mask that included all BAs in prefrontal non-motor areas, including associative and limbic cortices, as defined in a previous study that assessed cortico-subthalamic connectivity using the anatomy-functional YeB atlas. Two-sample *t*-tests were performed for each voxel, with 1000 permutations. *P*-values were corrected using familywise error (FWE) with the threshold-free cluster enhancement approach (TFCE), which could provide reliable correction at the voxel level (39).

## Results

### Demographic data

Mean demographics are provided in Table 1. The two PD patient groups and the HC group were comparable in age and cognitive performance, as assessed by the MoCA. The two PD patient groups were similar in disease duration and parkinsonian motor disability as assessed by the Movement Disorders Society-Unified Parkinson’s Disease Scale (MDS-UPDRS) part III OFF treatment (40). As expected, DBS patients had a significantly lower Levodopa equivalent daily dose (LEDD) than DOPA patients.

### Behavioral results

### Learning performances

All participants chose the most rewarded target in 80% of the trials for all sessions and reward probabilities (one-sample test compared to chance, p<0.001) in the last ten trials of the training phase (Supplementary Figure 2). PD patient’s performances did not significantly differ from HC’s (p>0.380). Learning performances were similar between ON and OFF sessions in the DOPA group (p>0.967) for both P=0.75 and P=1 blocks. DBS patients demonstrated significantly higher performance during OFF sessions (93%) compared to ON sessions (85%, p=0.035) for the P=0.75 block, while performances were comparable for the P=1 block (p=0.675).

**Figure 2.**
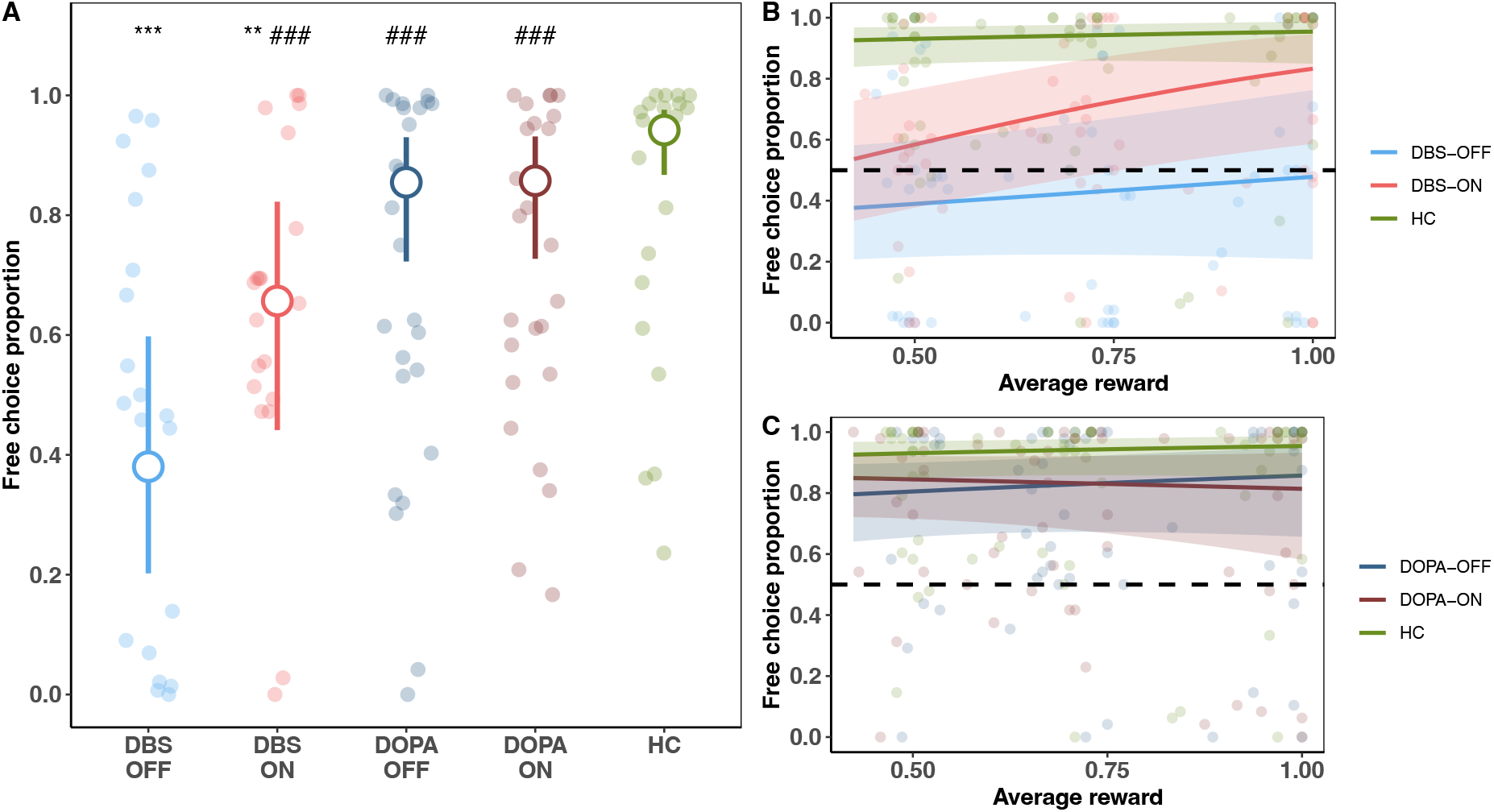
Free-choice preference at the first stage. (A) Free-choice preference. Small colored dots represent the mean individual free-choice preference. Large unfilled circles represent the estimated mean free-choice preference of participants. Error bars correspond to the 95% confidence interval (CI). (B-C) Free-choice preference as a function of the average obtained reward in the DBS (B) and DOPA (C) groups. Plain lines and ribbons represent linear estimates and the 95% CI, respectively. Colored dots represent mean individual performances. **: p<0.01, ***: p<0.001 compared to HC. ###: p<0.001 compared to DBS-OFF.

### Free-choice preference

Free-choice preference was higher than expected by chance for HC, and DOPA patients during ON and OFF sessions (estimated means [%] ± Standard Error (SE); HC: 94 ± 2 %, p<0.001; DOPA-ON: 86 ± 5 %, p<0.001; DOPA-OFF:85 ± 5 %, p < 0.001). Free-choice preference was in average comparable to the chance level in the DBS group, either OFF (38 ± 10 %, p = 0.279) or ON (66 ± 10 %, p = 0.151) sessions (figure 2A). Behavioral preferences were expressed from the onset of each reward probability block and tended to remain constant during the testing phase when looking at trial-by-trial preferences. (Supplementary Figure 3).

**Figure 3.**
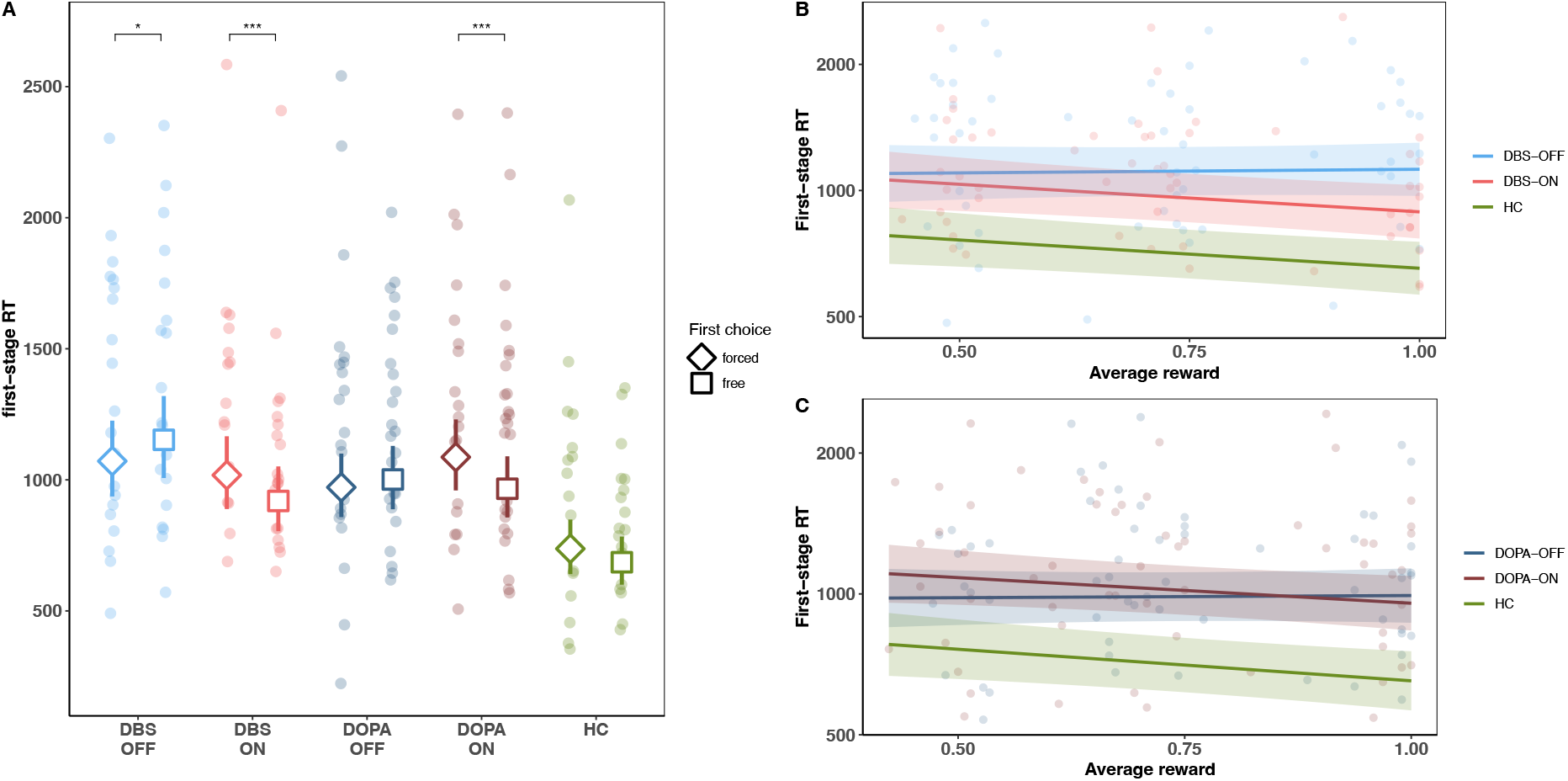
A: Reaction times (RT) during first-stage selection. Small colored dots represent the mean individual RTs. Large unfilled circles (*forced)* or squares (*free)* represent the estimated mean RT. B: RT estimated means during the first stage as a function of the obtained reward for the DBS and HC groups. C: RT estimated means during the first stage as a function of the obtained reward for the DOPA and HC groups. Error bars and ribbons correspond to the 95% CI.

#### Free-choice preference in the DBS group

Compared to HC, the preference for choice of DBS patients was lower during OFF (p<0.001) and ON (p = 0.005) sessions (Figure 2A). Importantly, DBS increased choice preference, as we observed lower levels of free-choice during OFF compared to ON sessions (p < 0.001). We found no main effect of the obtained reward (p = 0.349) and no interaction obtained reward x Session (p > 0.9). However, exploratory post-hoc analyses showed that the free-choice proportion increased with the obtained reward the during ON sessions (i.e. strongest free-choice proportion in block 3 where extrinsic reward delivery was deterministic, p = 0.003; Figure 2B).

Patients were also faster selecting *free* compared to *forced* choice during ON sessions (*free*: 919 ± 63 milliseconds [ms] vs. *forced*: 1018 ± 71 ms; p < 0.001), while patients were slower selecting *free* compared to *forced* choice during OFF sessions (*free*: 1153 ± 79 ms vs. *forced*: 1071 ± 73 ms; p = 0.021, figure 3A). When comparing ON and OFF sessions, patients were faster selecting *free-choice* options during ON than OFF (p < 0.001). There was no significant difference between ON and OFF sessions for *forced* choice options (p = 0.294), and no interaction obtained reward x *free/forced* choice options (p > 0.9). However, exploratory post-hoc analyses showed that patients responded faster with increasing obtained reward during ON (p = 0.016) but not OFF sessions (p = 0.861, figure 3B).

We found a similar pattern of reactions when analyzing patients’ behavior during the second stage of the task (i.e., when selecting rewarded targets, Supplementary Figure 4). ON-DBS patients were faster at selecting the second-stage target in *free* compared to *forced* trials (*free*: 830 ± 56 ms vs. *forced*: 891 ± 61 ms; p=0.021), while the opposite was observed during OFF-DBS sessions (*free*: 1059 ± 72 ms vs. *forced*: 960 ± 65 ms; p<0.001).

**Figure 4.**
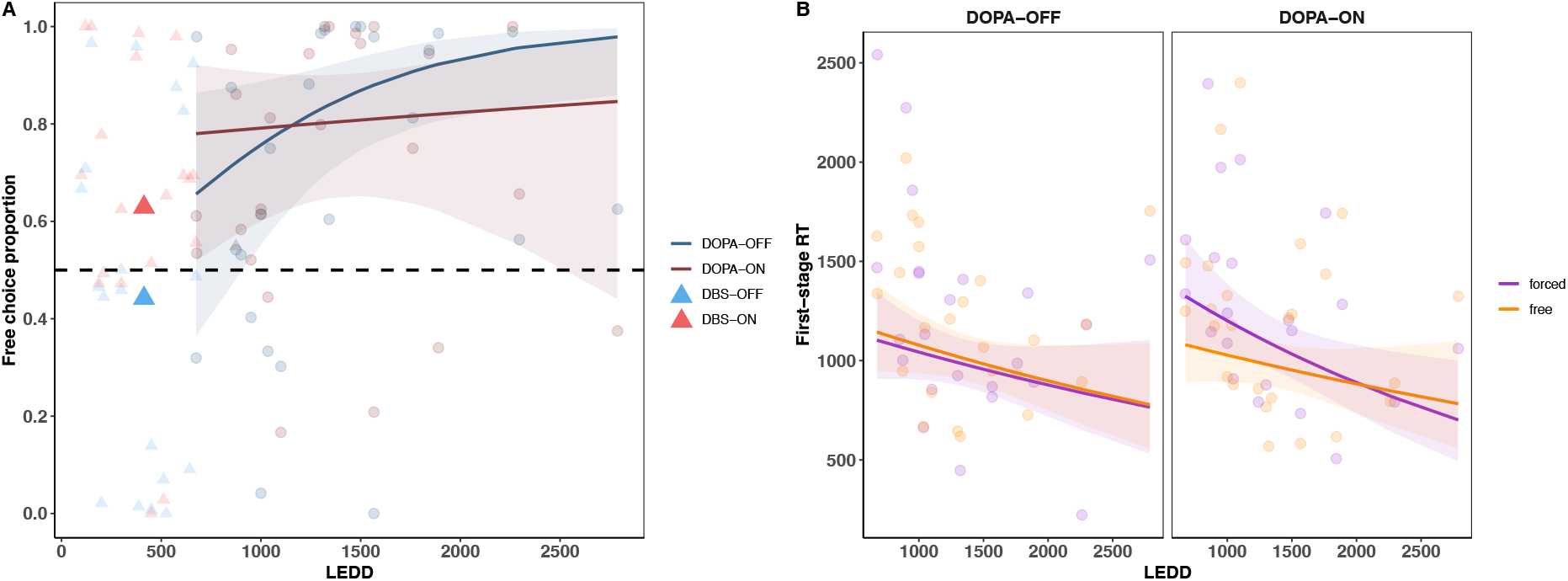
First stage free-choice preference of the DOPA group. (A) Free-choice preference plotted as a function of LEDD. Plain lines represent the linear estimates. Ribbons represent the 95% CI. Circles represent the preference of individual patients in the DOPA group. Large and small triangles are for visual comparison only and represent mean and individual free-choice preferences for DBS patients. Blue and red colors represent OFF and ON treatments, respectively. (B) ON and OFF first-stage RTs as a function of LEDD and first-stage choice. Violet (free) and orange (forced) dots represent the average RT at the individual level. Plain lines represent the linear estimates. Ribbons represent the 95% CI.

#### Free-choice preference in the DOPA group

Free-choice preference in DOPA patients was comparable to HC during OFF (p=0.436) and ON sessions (p=0.457) sessions (figure 2A). There were no significant differences between OFF and ON-DOPA sessions (p=0.999).

Patients were faster to select the *free* compared to the *forced* option in the ON-DOPA sessions (*free:* 966 ± 59 ms vs. *forced*: 1087 ± 69 ms, p<0.001), but not in the OFF sessions (*free*: 1001 ± 61 ms vs. *forced:* 971 ± 62 ms, p=1, figure 3A). Conversely, in the OFF-DOPA sessions, patients were slower to select the *forced* compared to the *free* option (p<0.001), with no significant differences during ON sessions (p=0.451). DOPA patients were also faster with increasing obtained reward during ON sessions (p=0.037), but not in the OFF sessions (p=0.861). We found no interaction of obtained with *free/forced* options (p>0.9, figure 3C). DOPA patients also selected the most rewarded target, with a higher proportion in all probability blocks (p<0.001), with no significant differences compared to HC (p>0.551).

During the second stage of the task, ON-DOPA patients were faster at selecting the second-stage target in *free* compared to *forced* trials (*free*: 871 ± 53 ms vs. *forced*: 1011 ± 63 ms; p<0.001). there was no significant *free* vs. *forced* difference during OFF sessions (*free*: 889 ± 54 ms vs. *forced*: 933 ± 58 ms; p= 0.610, Supplementary Figure 4).

#### Comparison of free-choice preference between DBS and DOPA patients

*Free-choice* preference was significantly lower for DBS-OFF patients compared to DOPA-ON (p<0.001) and DOPA-OFF (p<0.001).

#### Effect of chronic DA therapy on free-choice preference

Since we found no acute DOPA effect, we investigated whether differences in chronic DA therapy could have influenced choice preference (Figure 4). We considered the total amount of DA therapy (Levodopa Equivalent Daily Dose, LEDD) as a predictor of free-choice preference. We then included the LEDD as a continuous variable in the model and investigated its interactions with ON and OFF-DOPA sessions and the obtained reward. We found no main effect of LEDD (p=0.11) but found a significant LEDD x Session interaction (p<0.001, figure 4A). Higher LEDD influenced *free-choice* preference in DOPA patients during OFF sessions (p=0.024), while this result was non-significant during ON sessions (p=0.746). The comparison of LEDD slopes between ON and OFF-DOPA sessions was also significant (p<0.001). We found no significant interactions between LEDD x Obtained Reward (p=0.6) or LEDD x Obtained Reward x Session (p=0.8).

We also explored how LEDD influenced reaction times during the first stage. We built a subsequent model with LEDD, Session, *free/forced* choice, obtained reward and all their interactions as predictors for first-stage reaction times (figure 4B). We found no main effect of LEDD but a significant interaction LEDD x *free/forced* choice x Session (p=0.028). LEDD influenced reaction times for patients during ON-DOPA in *forced* trials (p=0.011), but not during ON-DOPA in *free* trials (p>0.182).

### Tractography results

We found that the connectivity between the right VATs and a subregion including the right medial prefrontal cortex (MPFC), pre-supplementary motor area (pre-SMA) and anterior cingulate cortex (total volume significant voxels: 436 mm^3^) was significantly higher in DBS patients who selected more the *free-choice* option ON compared to OFF-DBS session (Figure 5). This difference in connectivity was mainly explained by the differences between patients’ implantation sites, with therapeutic contacts located more anteriorly in the right compared to the left hemisphere (anteroposterior coordinates in the AC/PC referential; left: y = 10.2 mm; right: y = 10.8 mm; *W* = 431, p= 0.004, Supplementary Figure 5).

**Figure 5.**
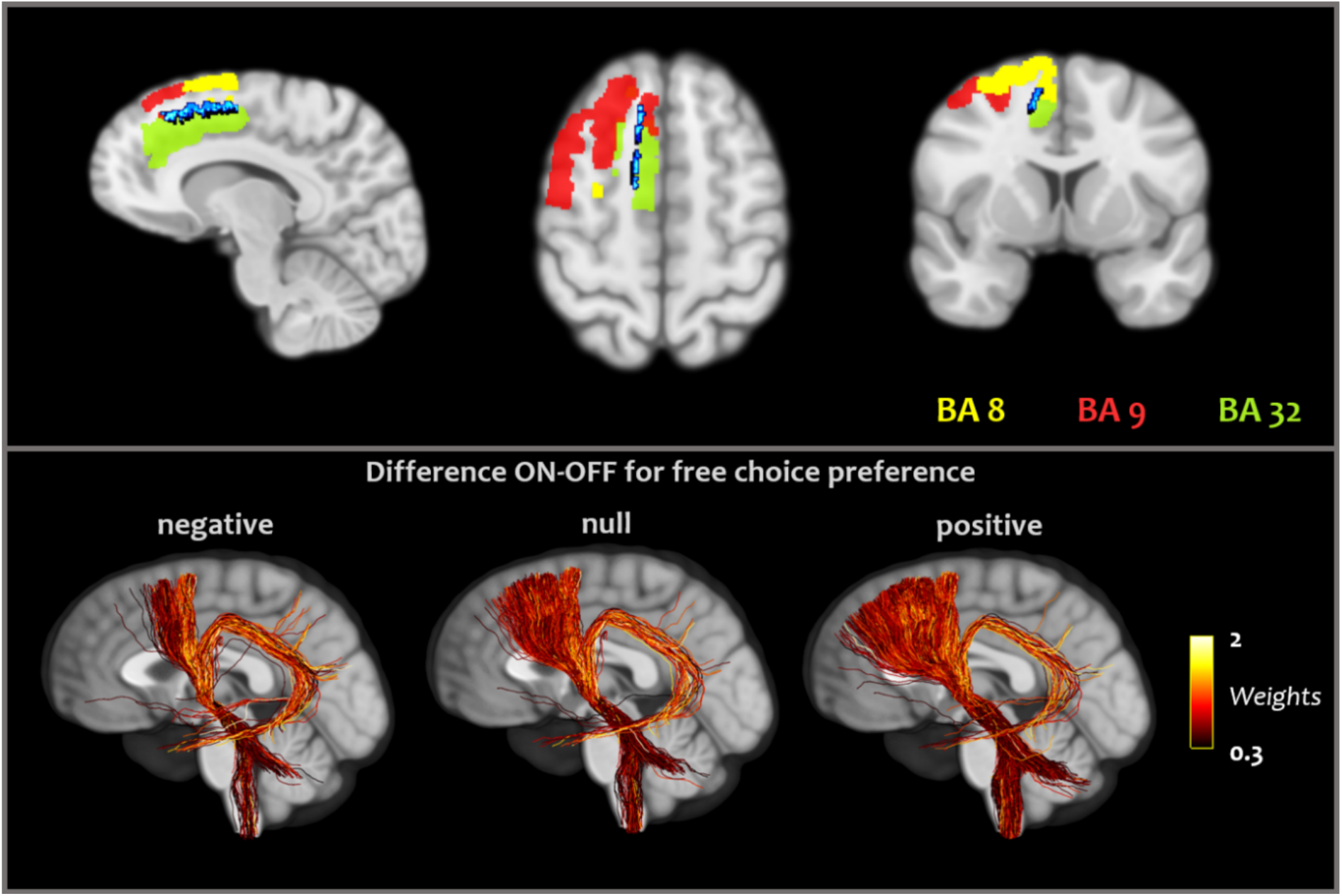
Voxelwise analysis between low and high ON-OFF free-choice preference in the DBS group. (Top) The blue region corresponds to the statistical map with a threshold of *P*<0.05 (FWER-TFCE) overlaid on a T1 volume of the template. Yellow, red and green regions correspond to the Brodmann areas 8, 9 and 32, respectively. (Bottom) Illustrations of connectivity in three patients with different effects of DBS on free-choice preference. Left to right represent streamline projections of individual patients with negative (−0.23), null and positive (0.97) effect of the stimulation on free-choice preference. The images are displayed using the radiological orientation (R: right).

To provide a visualization of the right prefrontal streamlines density distribution in the STN related to free-choice preference, we computed two T-maps, one corresponding to the mean streamlines density distribution activated by VATs within the STN (figure 6A) and the other the T-values of the mean difference of streamlines density distribution between the two groups based on their ON-OFF differences for *free* option selection (Figure 6B). We observed that the maximal activated streamlines density in patients with high ON-DBS free-choice preference compared to OFF-DBS was located in a more anterior position within the STN (anteroposterior coordinates in the AC/PC referential; y=11 mm vs. y=9 mm, Figure 6C). The right VATs also tended to be larger than the left VATs, but this was not statistically significant (mean VAT for the right and left STN= 95 and 74 mm^3^, respectively, p= 0.231).

**Figure 6.**
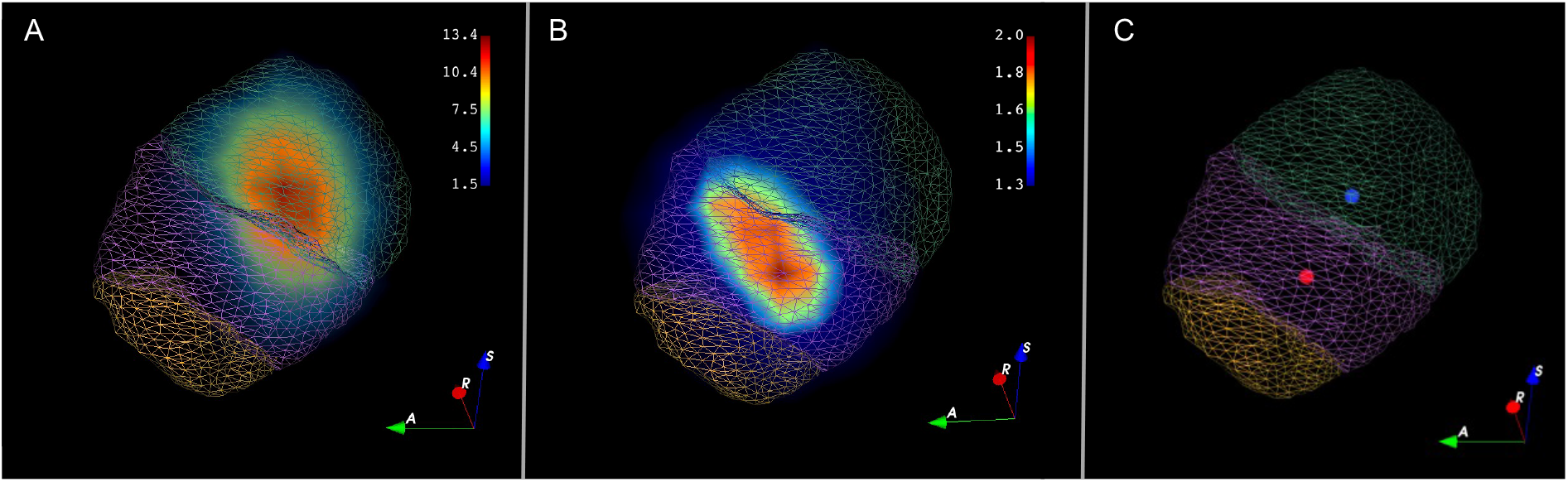
T-maps of mean streamlines density distribution overlapping with the VATs in the whole right STN (A) and the mean difference between patients with a DBS choice effect (B, median-split on the ON minus OFF free-choice selection). T-maps are superimposed on the STN limbic (yellow), associative (purple) and sensorimotor (green) schematic anatomo-functional territories from the Yeb atlas (41). Colorscales represent the T-values. A, R, and S for anterior, right and superior, respectively. (C) Maximal activated streamlines density in the right STN overlapping with the VATs (blue dot) and the mean difference between patients with a DBS choice (median-split on the ON minus OFF free choice selection, red dot). Coordinates in millimeters in the AC/PC referential system, relative to the posterior commissure: x=13.71 mm; y=9 mm; z=2 mm (blue dot) vs. x= 11 mm; x=11 mm; z=3 mm (red dot).

## Discussion

Intrinsic motivation accounts for many human behaviors independent of extrinsic outcomes (3). Neural mechanisms underlying intrinsic motivation remain unclear, and no direct evidence showed that DA and basal ganglia systems could be involved in humans. Here, we used choice opportunities as an intrinsic reward and tested patients with PD. First, we showed that STN-DBS enhances free-choice preference, and this effect depends on a MPFC-STN network. We also found that levels of chronic DA therapy positively correlated with free-choice preference. These findings suggest that deep hypodopaminergic states such as chronic low levels of DA therapy diminished the preference for choice opportunities, and this preference can be re-established either with high doses of daily DA therapy or STN-DBS with direct modulation of right MPFC projections.

### STN DBS increases preference for risky choices

We observed that OFF-DBS patients displayed no difference between the free and forced choice options, whereas ON-DBS patients showed an increased preference for free-choice. This improvement was linked to enhanced connectivity between the right MPFC and individual VATs. The MPFC-STN network, which is part of the hyperdirect pathway, has been classically related to processing conflicting information and response inhibition, which could be modulated directly by DBS (23,28,31,42). Therefore, PD patients exhibiting increased post-DBS connectivity between these regions demonstrate impulsiveness in both behavioral tasks and neuropsychiatric evaluations (24,28,43). Computational modeling suggests that STN-DBS could create a more permissive decision threshold (22,23), particularly in conflict-laden situations that result in ambiguous or irrational decisions where the ratios of gains to losses are not easily discernible, thereby posing a risk of losing rewards. In our experiment, STN-DBS resulted in a faster and enhanced preference for free-choice. This increase in choice preference was especially pronounced in blocks with high reward probabilities (i.e., P>0.5), where opting to choose could potentially lead to reward losses. In this context, such risky behavior contradicts the economically rational decision, which would have been to select the safer option with no choice opportunity. Consequently, ON-DBS patients exhibit a risky attitude by over-selecting the free option, particularly in conditions where reward probabilities are not stochastic. STN-DBS, by disrupting the inhibitory responses mediated by the right MPFC-STN pathway, may allow participants to express their preference for immediate intrinsic rewards in the initial phase of the task while disregarding the potential loss of an extrinsic reward in the subsequent phase. Consistent with previous findings, this result implies that the intrinsic rewarding aspect of choice may, at least in part, stem from its riskier nature (8). Here, making a risky decision can be analyzed in a way that enhances the agent’s perception of the associated action and significantly impacts the environment. This realization fosters a sense of agency and reinforces the perception of one’s ability to exert control over one’s life (8,44). Thus, the relationship between the appropriate MPFC activity and free-choice preference may be linked to voluntary actions to control the environment. The MPFC is indeed part of a network that computes action-outcome associations, enabling more flexible motor plans to achieve internal goals (45). This function is supported by the fact that impairments in this area, as well as pre-SMA dysfunctions in PD, are connected to deficits in flexible behaviors that could be alleviated by STN-DBS (46–48). In our study, OFF-DBS patients expressed no preference between free and forced options, suggesting that STN-DBS may promote the emergence of free-choice behaviors as a self-driven incentive by restoring a flexible decision-making process.

However, while these results indicate that modulation of the hyperdirect pathway can enhance intrinsic motivation for choice opportunities by modulating response inhibition, the role of DA in promoting a preference for intrinsic rewards is also further indicated by the behaviors of DBS patients.

### Low dose of DA therapy diminished free-choice preferences

In the OFF-DBS patients, we observed the lowest preference for *free-choice* that could be a consequence of the most substantial reduction of DA medication in this group. Moreover, we found that patients in the DOPA group preferred free-choice options independently of the acute levodopa challenge. We found a positive association between the LEDD and the preference for free-choice in OFF-DOPA patients. We hypothesized that chronic instead of acute DA therapy may affect more significantly behavioral responses. The long-term effects of DA therapy on enhancing reward-based values of appetitive stimuli and reducing sensibility to aversive outcomes have been constantly reported in impulsive and compulsive behaviors in which the incidence is correlated to the long-term exposition of DA therapy (49–51). The mechanism underlying such enhanced appetitive behavior depending on chronic dopamine treatment remains poorly understood. Chronic DA impregnation could influence synaptic regulation of DA transmission, down-regulation of D2 auto-receptors, and enhanced spontaneous activity of DA neurons, thus potentiating DA phasic responses induced by rewarding stimuli (49,52). Another explanation could be levodopa’s long-duration response (LDR). LDR is observed in patients who maintained a reminiscent pharmacological motor improvement for more than several hours (or days) after levodopa withdrawal (52,53). Although this phenomenon is under-studied, it has been hypothesized that long-term effects could account for more than 60% of the motor benefit in PD patients (53). LDR is even more under-studied regarding non-motor symptoms, but evidence suggests its involvement in motor learning and motivational vigor (53,54).

Regarding the relation between DA and intrinsic motivation, this result sustains the hypothesis that the same network encodes both *intrinsic* and *extrinsic* rewards. Free-choices may act as a *premium* or *bonus* when selecting rewarding items, thus enhancing their subjective value and reinforcing free-choice options in the long term. Such *bonus* has been previously modeled in reinforcement learning paradigms in healthy humans (55–57). It has also been recorded through specific phasic midbrain DA activity related to intrinsic rewards in an electrophysiological study on non-human primates (13). DARPP32 polymorphism, a gene associated with DA signal transduction in the basal ganglia, is also linked to an increased preference for freely chosen over forcedly sampled items (55). Following this hypothesis, enhanced free-choice preference due to chronic DA exposure may relate to the potentiation of appetitive signals related to *intrinsic* rewards in the same way as *extrinsic* ones.

### Limitations

We did not find an acute effect of DA therapy on free-choice proportions in the DOPA group when comparing ON and OFF states. However, similar to DBS patients, ON-DOPA patients were faster in selecting free vs. forced options, potentially reflecting motivational processes. The lack of difference in free-choice proportions might stem from our population’s characteristics of advanced patients with PD. These characteristics include prolonged exposure to high-dose DA therapy, which may limit behavioral changes after a 12-hour cessation.

Additionally, task performance order raises the question of test-retest effects, but no significant differences were found between successive HC sessions. However, we observed intra-session differences in reaction times between *free* and *forced* options, and learning performance showed no significant ON-OFF differences for DOPA patients. In contrast, DBS patients performed better OFF than ON, with results above 80% of good answers for both sessions.

Finally, we used a normative connectome instead of individual acquisitions for tractography analysis. While this approach may minimize signal-to-noise ratios, it also reduces individual variability. Notably, this approach has been found to yield comparable results to individual acquisitions in patients with PD (58). Furthermore, our connectome was derived from a population closely matching our study cohort.

## Conclusion

Our study demonstrates that dopamine depletion diminishes intrinsic motivation for making choices, particularly in risky scenarios. However, this can be restored to normal levels by modulating the prefrontal-subthalamic pathway via deep brain stimulation or chronic DA therapy. Future research should explore whether intrinsic motivation and extrinsic reward processing deficits underlie broader motivational deficits in patients.

## Acknowledgments

D.B. was supported by a FRM fellowship (FDM201906008526). C.K. and B.L. were supported by an Agence Nationale de la Recherche (ANR) grant, ANR-19-CE37-0014-01. J.M. was supported by the ANR grant ANR-19-CE37-0014-01 (ANR PRC) and by the European Commission (H2020-MSCA-IF-2018-#845176). The authors are grateful to the staff of the neurology and neurosurgery departments of the Pitié Salpêtrière Hospital for their excellent patient care during the investigation. Anonymized data used for the analyses are available upon reasonable request.

