## Supplementary information for "Subthalamic deep brain stimulation and chronic dopaminergic therapy enhance free-choice seeking in patients with Parkinson’s disease"

### Supplementary methods

#### Inclusion criteria

Inclusion criteria for PD patients were a diagnosis of Parkinson's disease following the United Kingdom Parkinson's Disease Society Brain Bank clinical diagnostic criteria (1). Patients without significant cognitive impairment (Montreal cognitive assessment [MoCA] ≤ 25/30) or concomitant psychiatric disease were included. Inclusion criteria for healthy controls were the absence of neurological or psychiatric disorders and the absence of significant cognitive impairment (MoCA ≤ 27/30). All the participants were mandated to exhibit a good understanding of the task instructions

#### Image processing

Preoperative MRI volumetric T1-weighted, T2-weighted FLAIR (fluid attenuated inversion recovery) sequences and postoperative computerized tomography (CT) scans were obtained for the 20 patients of the DBS group. We pre-processed images with LEAD-DBS pipelines using the default settings (2).

CT images were co-aligned to MRI T1-weighted sequences using a two-stage linear alignment as implemented in Advanced Normalization Tools (3), and the MRI FLAIR sequence was linearly co-aligned to the T1-weighted sequence using SPM12 (4). Pre- and post-operative acquisitions were spatially normalised into MNI_ICBM_2009b_NLIN_ASYM space (5) using the SyN alignment approach as implemented in Advanced Normalisation Tools (3), adding two supplementary nonlinear SyN-alignments that consecutively focused on the basal ganglia as defined by subcortical masks in Schönecker et al. (6). Visual verification and manual correction, when necessary, were performed at every stage of the process.

DBS electrode localisation was corrected for brain movement in postoperative acquisitions by applying a refined affine transform calculated between pre- and post-operative acquisitions that were restricted to a subcortical area of interest, as implemented in the brainshift-correction module of Lead-DBS software (7). Electrodes were automatically pre-localised using the Precise and Convenient Electrode Reconstruction (PaCER) algorithm (8). For one patient, the PaCER algorithm failed and the trajectory reconstruction and contact reconstruction (TRAC/CORE) was used instead (7). Visual verification of the reconstruction and manual correction were performed if necessary. For patients with directional leads, orientation was determined using the Direction orientation detection (DiODE) algorithm implemented in LEAD-DBS (9). Using LEAD-DBS (10), coordinates were then converted from the MNI space into anterior commissure and posterior commissure(AC/PC) coordinates (relative to the posterior commissure point).

After normalising and localising electrode trajectories, the VAT for each electrode was estimated using a finite element approach based on a four-compartment tetrahedral mesh, as detailed in Horn et al (11). The resulting estimated E-field was thresholded above 0.2 V/mm to obtain finite volumes.

### Supplementary figures

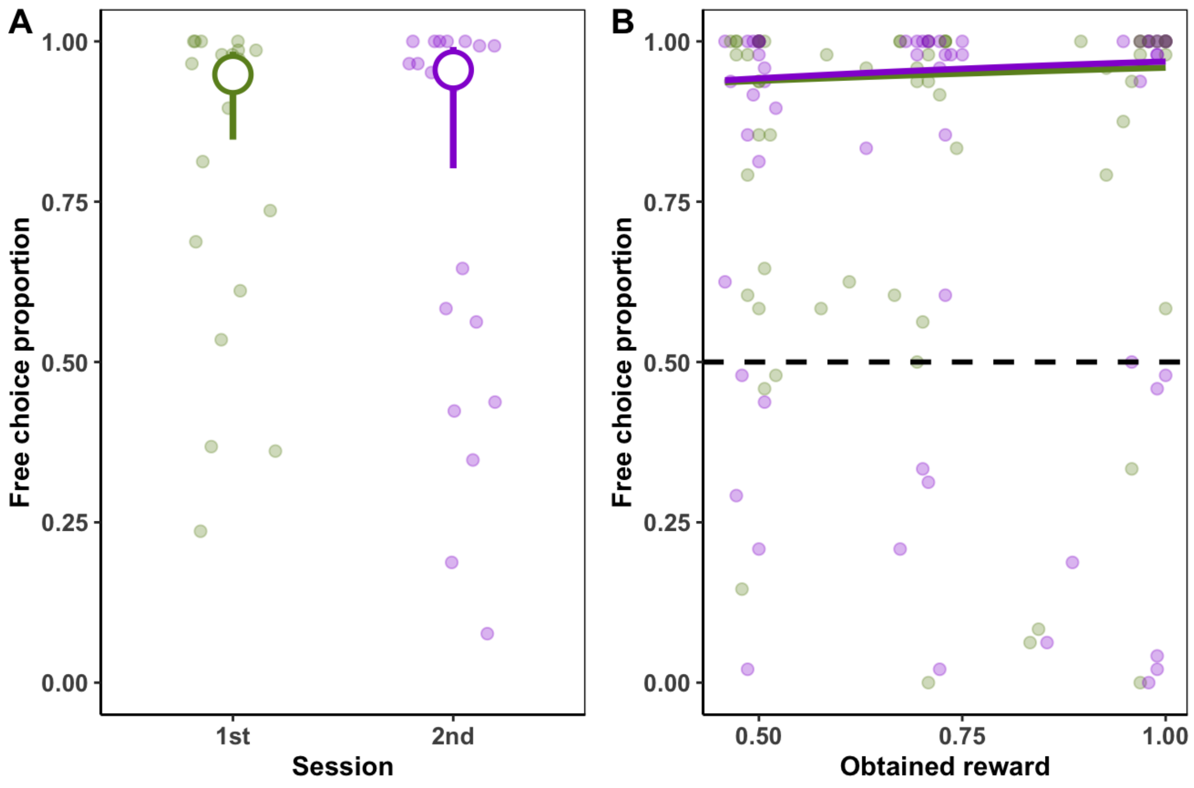


**Supplementary Figure 1. Free choice preference of healthy controls during two consecutive sessions. A.** Mean free choice proportions between the first and second sessions. Colored circles are the mean estimates, and error bars correspond to CI 95% intervals. We found no significant effect for session order in this group (p=0.707) **B.** Free choice proportions plotted against the average obtained reward. We found no differential effect of reward probability between the first and second session (p= 0.792). Plain lines are the mean estimates. Colored dots express individual means values.

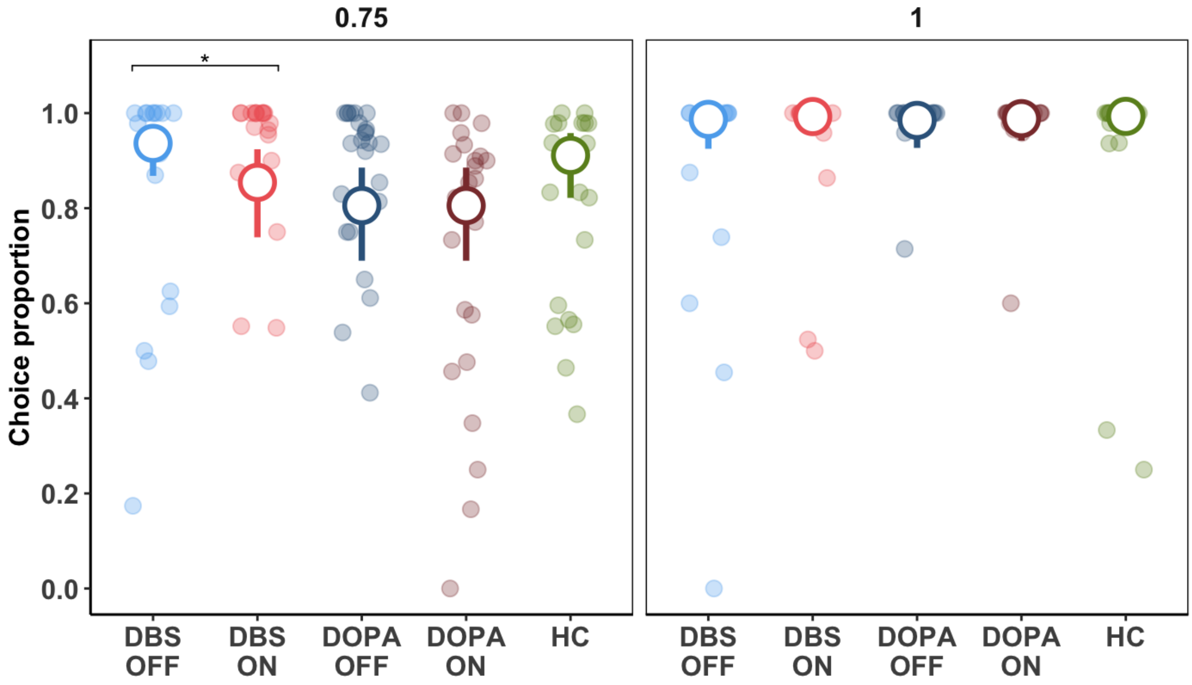


**Supplementary Figure 2. Learning performances at the end of the training phase (last ten trials)**. Large unfilled circles represent the estimated means. Error bars correspond to the 95% confidence interval (CI). Colored dots represent mean individual performances. *: p<0.05

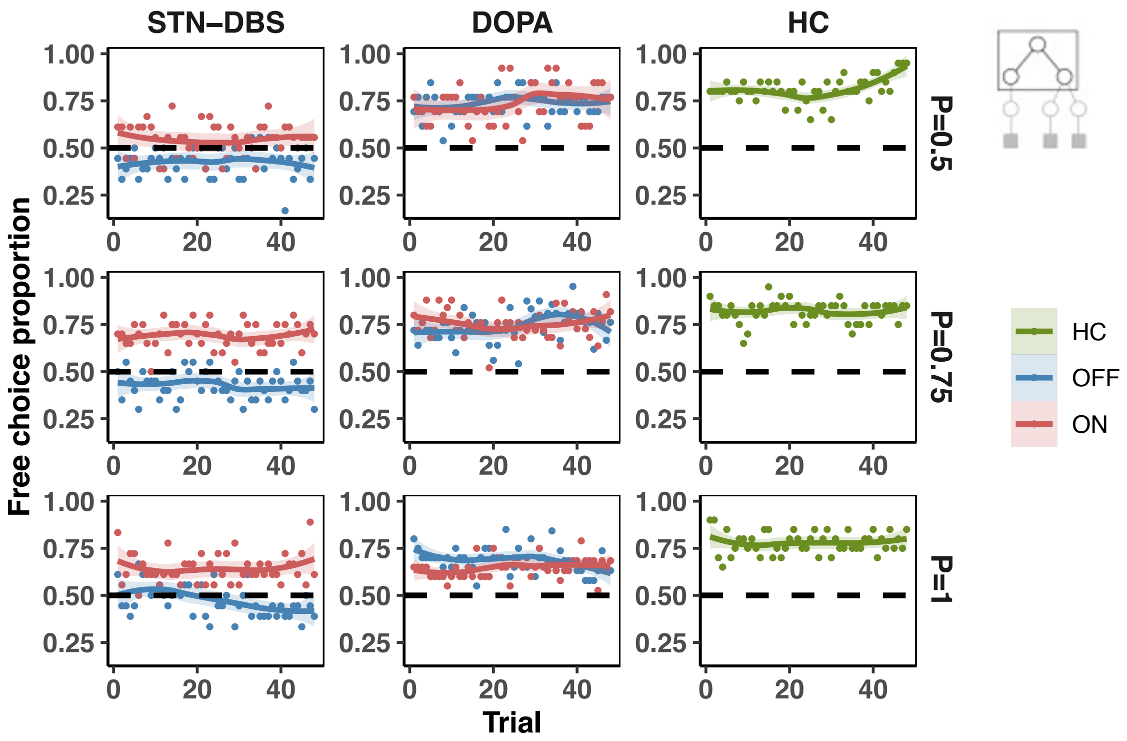


**Supplementary Figure 3. Temporal dynamic of the selection of *free* choice preference at the first stage across the task per block**. Small dots represent mean free-choice preferences at the trial level among all patients. Lines represent LOESS smoothing along with their respective standard errors. The horizontal dashed lines represent chance level. Based on linear regressions (free choice proportion ~ trial), there were no gradual changes in preference for choice throughout the trials ($p > 0.1$), except for OFF-DBS patients during the P=1 block (trend = -0.043, $p = 0.001$).

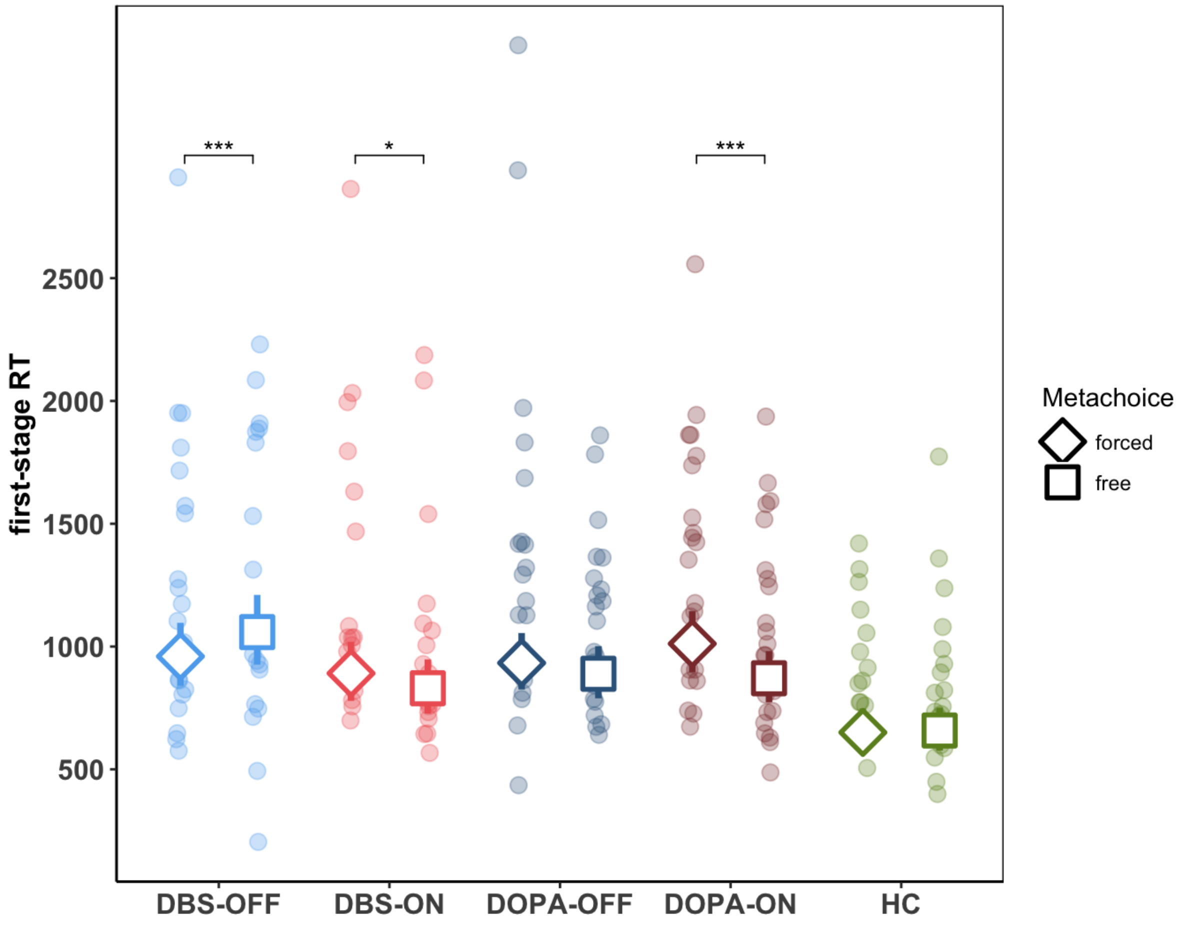


**Supplementary Figure 4. Reaction times (RT) during second-stage selection.** Small colored dots represent the mean individual reaction times (RTs). Large unfilled circles (forced) or squares (free) indicate the estimated mean RT. Error bars reflect the 95% confidence interval. *: $p<0.05$, ***: $p<0.001$

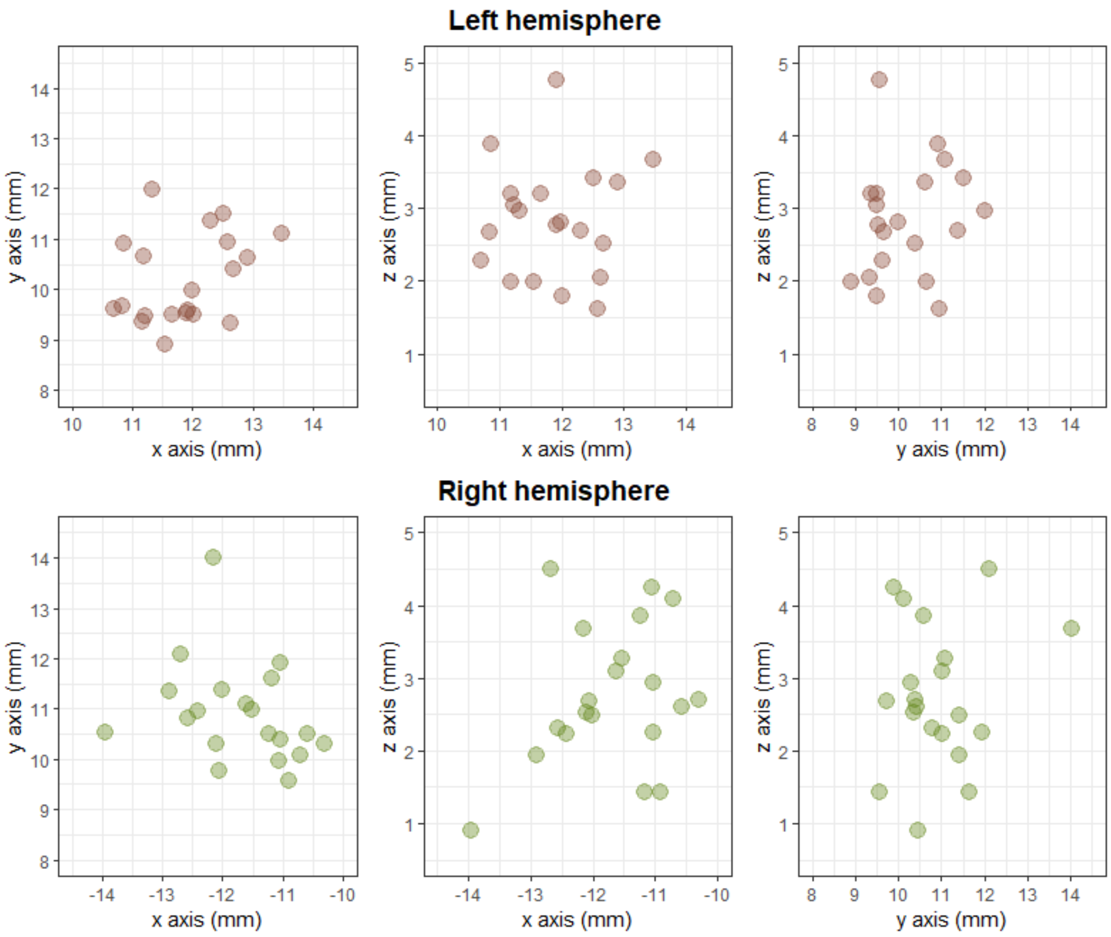


**Supplementary Figure 5. Individual variability of active contacts coordinates.** Each point represents one patient's active contact in the x (lateral), y (antero-posterior), z (depth) axis. Coordinates are given in millimeters in the AC/PC referential system, relative to the posterior commissure.

### Supplementary Table 1 Stimulation parameter settings for each patient.

| Patient ID | Manufacturer | Left hemisphere: Active contact(s) | Left hemisphere: Amplitude (mA or V)[a] | Left hemisphere: Pulse duration (μs) | Left hemisphere: Frequency (Hz) | Right hemisphere: Active contact(s) | Right hemisphere: Power (mA or V) | Right hemisphere: Pulse duration (μs) | Right hemisphere: Frequency (Hz) |
|---|---|---|---|---|---|---|---|---|---|
| 1 | Boston | 5(75%)-7(25%) [b] | 2,70 | 60 | 130 | 4 | 4,00 | 60 | 130 |
| 2 | Boston | 5 | 3,50 | 100 | 130 | 6 | 2,90 | 100 | 130 |
| 3 | Medtronic | 2 | 2,60 | 60 | 90 | 2 | 3,10 | 60 | 90 |
| 4 | Medtronic | 2 | 2,00 | 60 | 130 | 1 | 2,20 | 60 | 130 |
| 5 | Medtronic | 1 | 3,40 | 60 | 130 | 2 | 3,10 | 60 | 130 |
| 6 | Boston | 5 6 7 | 2,90 | 60 | 130 | 5 | 2,80 | 60 | 130 |
| 7 | Medtronic | 1 | 1,80 | 60 | 130 | 2 | 3,00 | 60 | 130 |
| 8 | Boston | 5 6 7 | 3,80 | 60 | 130 | 2 3 4 | 3,40 | 60 | 130 |
| 9 | Boston | 2 3 4 | 3,60 | 60 | 130 | 2 3 4 | 2,60 | 60 | 130 |
| 10 | Boston | 2 3 4 | 2,80 | 60 | 130 | 2 3 4 | 3,80 | 60 | 130 |
| 11 | Boston | 5(75%)-6(25%) [b] | 3,60 | 60 | 130 | 5(2%)-6(8%)-3(68%)-2(22%) [b] | 5,00 | 90 | 130 |
| 12 | Boston | 5 6 7 | 1,40 | 60 | 130 | 2 3 4 | 2,10 | 70 | 130 |
| 13 | Boston | 5 6 7 | 2,60 | 60 | 130 | 2 3 4 | 2,40 | 60 | 130 |
| 14 | Boston | 3 | 1,70 | 60 | 130 | 4 | 2,10 | 60 | 112 |
| 15 | Medtronic | 2 | 2,35 | 60 | 130 | 3 | 3,35 | 60 | 200 |
| 16 | Medtronic | 2 | 2,70 | 60 | 130 | 1 | 3,40 | 60 | 130 |
| 17 | Medtronic | 2 | 2,40 | 60 | 130 | 2 | 3,20 | 60 | 130 |

| | | | | | | | | | |
|---|---|---|---|---|---|---|---|---|---|
| 18 | Boston | 5 6 7 | 3,80 | 60 | 179 | 8 (20%)<br>6(40%)<br>5(40%) [b] | 3,90 | 60 | 179 |
| 19 | Medtronic | 2 | 3,70 | 60 | 130 | 2 | 2,80 | 60 | 130 |
| 20 | Medtronic | 2 | 2,70 | 60 | 205 | 2 | 1,85 | 60 | 195 |

[a] Amplitude of stimulation are given in milliamperes (mA) for Boston Vercise Cartesia and volts (V) for Medtronic 3389 electrodes.

[b] Percentages between brackets correspond to the proportion of current delivered for each active contact.
